# First Sin Nombre virus (*Orthohantavirus sinnombreense*) genome sequences from the Northwestern United States

**DOI:** 10.1101/2025.09.10.675434

**Authors:** Grant Rickard, Ricardo Rivero, A. Catherine Grady, Jennifer Horton, Cody J. Lauritsen, Stephen Fawcett, Samuel M. Goodfellow, Hanna N. Oltean, M. Pilar Fernandez, Stephanie N. Seifert

## Abstract

We report the first Sin Nombre virus (SNV) genome sequences from the Northwestern United States and the first SNV sequences recovered from voles. Analysis of samples collected from 189 individual rodents revealed high SNV prevalence in the region and evidence of viral reassortment, highlighting ongoing viral diversification in rodents.

## Background

*Orthohantavirus sinnombreense* (Sin Nombre Virus, SNV) is a member of the genus *Orthohantavirus* (family Hantaviridae) and the primary cause of Hantavirus Pulmonary Syndrome (HPS) in North America. First identified during a 1993 outbreak in the Four Corners region of the United States, SNV is linked to severe respiratory disease and high mortality(1). As of 2024, more than 850 HPS cases have been reported in the United States, with a 36% case-fatality rate(2,3).

SNV is primarily maintained by *Peromyscus* spp. (deer mice), widespread rodents that are frequently associated with agricultural and peridomestic settings. Human infection usually results from inhalation of aerosolized viral particles from contaminated excreta(4,5), with zoonotic risk influenced by ecological factors affecting rodent density(6). While SNV is commonly detected in deer mice, several reported detections in sympatric rodent taxa and broad geographic distributions suggest greater complexity in virus maintenance(7,8).

Viral genomic surveillance can provide insights on viral evolution and spread(9), yet there are fewer than 100 full SNV genomes published and none from the Northwestern United States(10). Here, we report the first detection and genome sequences of SNV in the Northwestern United States and in voles.

### The Study

Between June 2023 and August 2023, we live-trapped rodents using Sherman traps at farms and natural areas in the Palouse region of eastern Washington and western Idaho (Figure 1), a major agricultural hub dominated by wheat and canola fields in the inland Northwestern United States. Sampling was conducted over three consecutive nights, providing repeated fecal sampling with mark-recapture, and lethal sample collection including tissue sampling on the third night. We collected and tested samples from 189 individual rodents across agricultural and natural landscapes in the Palouse. Sherman live traps were deployed at 8 farms and 2 forested sites in three 100 m X 30 m grids at each location. All captured rodents were identified, body and weight measurements collected, and either marked with an ear tag and released or euthanized following AVMA guidelines as approved by the Institutional Animal Care and Use Committee of Washington State University. Samples were collected under Idaho scientific collections permits #36112 and JOE19, and Washington state scientific collections permit SEIFERT 23-122.

**Figure 1.**
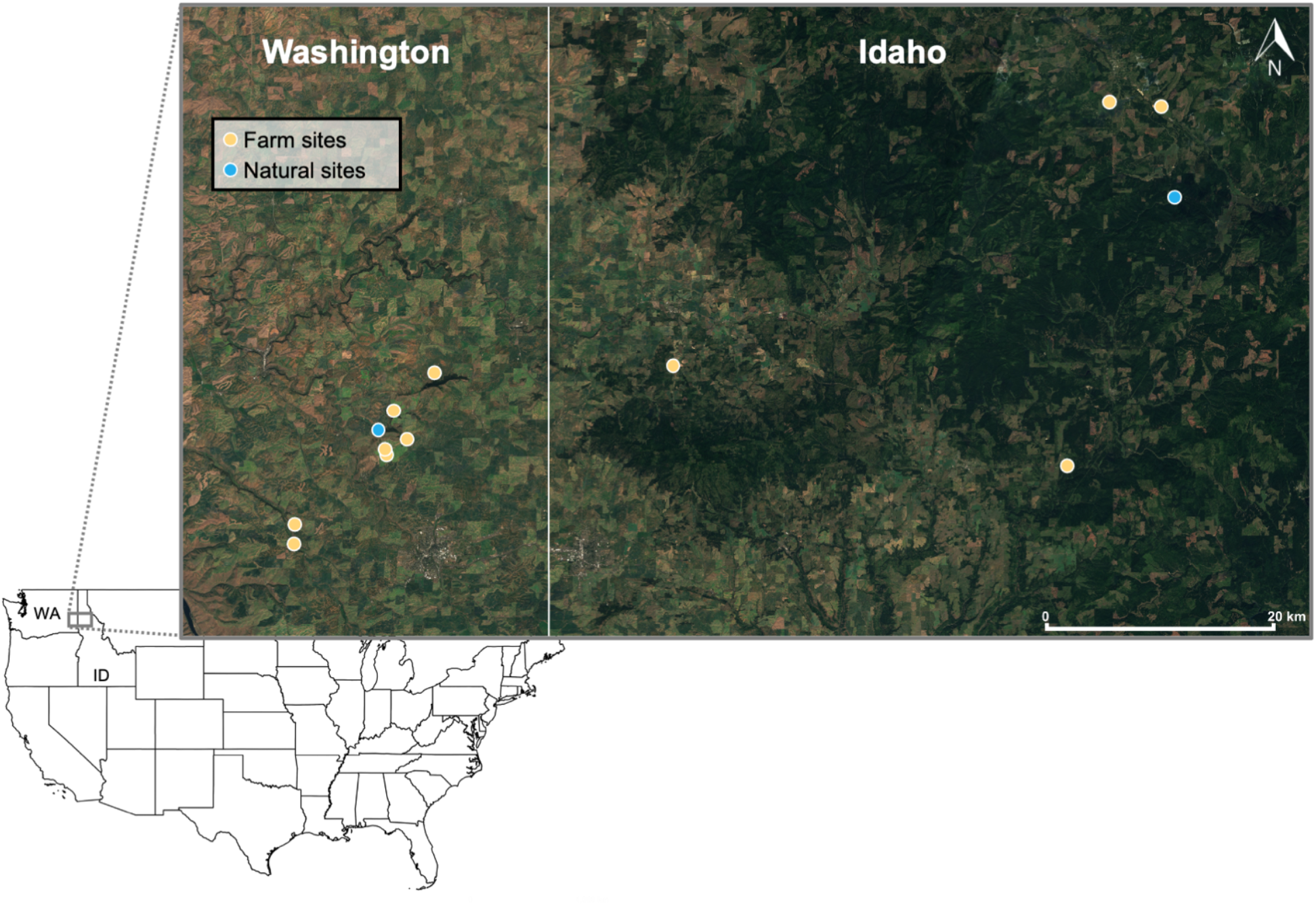
Locations of successful unique rodent sampling grids by land type; farms in yellow and natural areas in blue. Basemap is Sentinel-2 cloudless, EOX IT Services GmbH (CC BY 4.0). State boundary is U.S. Census Bureau Cartographic Boundary files (5m, public domain).

We collected fecal samples on all trapping nights along with serum, lung, and bladder tissue on the third and final trapping night at each site. As no fecal samples tested positive for SNV by RT-qPCR on mark-recapture days, we report only data from the final capture for each individual rodent sampled. Across all sites, we collected samples from 2 creeping voles (*Microtus oregoni*), 18 montane voles (*Microtus montanus*), 4 western meadow voles (*Microtus drummondii*), 2 house mice (*Mus musculus*), 153 western deer mice (*Peromyscus sonoriensis*), 1 yellow pine chipmunk (*Neotamias amoenus*), and 9 least chipmunks (*Neotamias minimus*).

We used recombinant nucleocapsid (N) protein from Sin Nombre virus (strain SN77734; BEI Resources, Manassas, Virginia, USA; NR-9670) containing a C-terminal histidine tag to assess seroreactivity in rodent serum samples. Briefly, the N protein was immobilized on Maxisorp-coated plates (ThermoFisher Scientific, Waltham, Massachusetts, USA; Cat# 439454), after which rodent serum diluted at 1:100 was applied. N protein bound antibodies were detected with horseradish peroxidase-conjugated goat anti-rat secondary antibody (AbCAM, Eugene, Oregon, USA; Cat #ab97057). Positive and negative thresholds for each plate were determined as the mean optical density of naïve serum controls plus 3 SD.

Total RNA was extracted from tissue samples and fecal samples using the Zymo Quick-RNA MagBead kit (Zymo Research, Irvine, California, USA; Cat #R2132) according to the manufacturer’s protocol for each sample type. Presence of viral RNA was detected by RT-qPCR as described in Williamson *et al*. 2021(11) with an internal control targeting the *Peromyscus maniculatus* hypoxanthine-guanine phosphoribosyltransferase (HPRT1) gene with in-house designed forward primer (5’-CAAAGCCTAAGAGGAGAGTTCA-3’), reverse primer (5’-GATGGCCGCAGAACTAGAA-3’), and probe (5HEX/AGGAGTCCC/ZEN/ATTGATGTTGCCAGT/3IABkFQ).

Unexpectedly, montane voles captured on farmlands showed the highest prevalence overall, with 50% seroprevalence and 22.2% RT-qPCR positive lung tissue samples (Table 1). Western deer mice tested positive by serology and RT-qPCR with higher prevalence on farms compared to natural areas (29.3% vs 20.0% seropositive, 9.4% vs 7.6% RT-qPCR positive lung tissue samples) and were the only species with an RT-qPCR positive fecal sample and bladder sample, in both cases from adult males (Table 1).

**Table 1.**
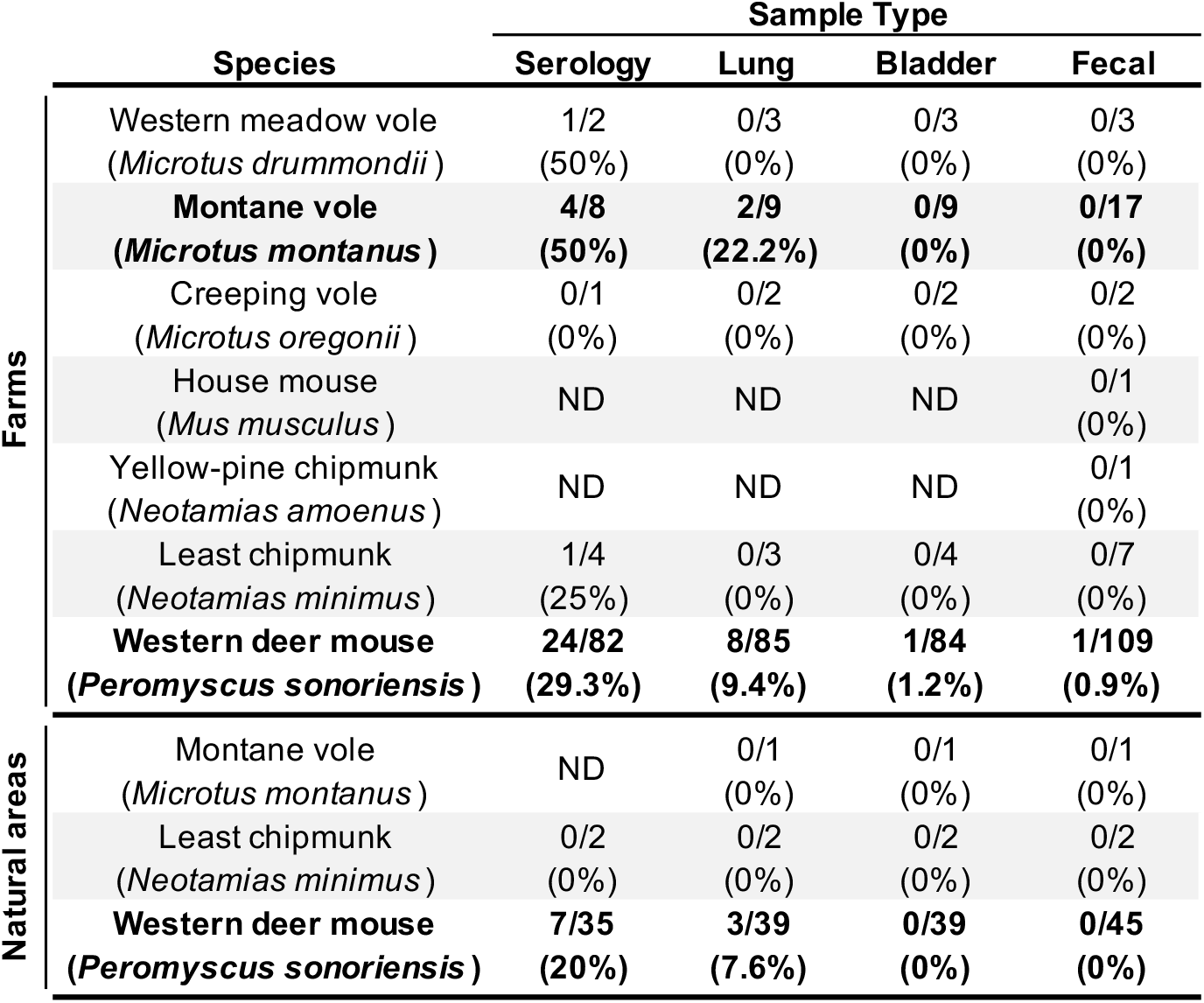
Serology and RT-qPCR results for Sin Nombre virus in rodent species sampled from farms and natural areas in the Palouse region of eastern Washington and western Idaho. Numbers indicate positives/total tested, with prevalence (%) in parentheses. Taxa with RT-qPCR positive samples in bold. ND = not done.

We modeled SNV lung positivity by RT-qPCR as a binary outcome using logistic regression with mean bias reduction (brglm2 package in R v4.1.3) for species with at least one RT-qPCR positive lung tissue sample. Predictors were land type (Forest or Farm), sex, age (Adult or Juvenile), and species (*Microtus montanus* or *Peromyscus sonoriensis*). Males had significantly higher odds of RT-qPCR positivity in the lung tissue samples than females (odds ratio [OR] = 6.27, confidence interval [CI] = 1.3 - 30.1), whereas land type, species, and age showed smaller effect sizes. Elevated SNV prevalence in male deer mice aligns with known SNV ecology, though the precise mechanism is unknown(12) and warrants further study.

We sequenced SNV using a tiled amplicon scheme proposed by Goodfellow *et al*.(13) on the Oxford Nanopore platform. Reads were quality-controlled, primers trimmed, reads mapped, and the consensus sequence extracted with an in-house assembly pipeline (https://github.com/viralemergence/SNVler). We recovered sequence data for all three SNV genome segments from 10 individual rodents including, for the first time, two montane voles with segment completeness ranging from 24-100% and depth between 6.8X-476.1X (GenBank Accession numbers TBD). To address persistent M-segment dropouts, we designed flanking primers (MsegFor 5’-GCAGGTAGCTGATCTCAAG-3’, and MsegR 5’-CCAGTCCATGTAAGAGGTAC-3’) for amplification and sequencing, improving assemblies and informing future refinement of the primer set for the Northwest. These are the first SNV genomes from the Northwestern United States, filling a critical knowledge gap(2).

Segment-wise alignments curated from NCBI Virus were analyzed in BEAST v1.10.5 (strict clock, exponential coalescent) with uniform tip sampling for incomplete collection dates(14). Estimated clock rates were 1.19×10^−4^, 1.203×10^−4^, and 1.223×10^−4^ subs/site/year, with root-to-tip regression resulting in R-squared values of 0.197 (p-value: 4.16e-04) for the S-segment, 0.067 (p-value: 0.0282) for the M-segment, and 0.257 (p-value: 1.056e-04) for the L-segment, suggesting a weak temporal signal. This likely reflects geographic population structure, reinforcing the need for broader sequencing to temporally resolve the SNV phylogeny. All Palouse sequences formed a clade closest to genomes from Montana collected in 2008/2009 (Figure 2)(11).

**Figure 2.**
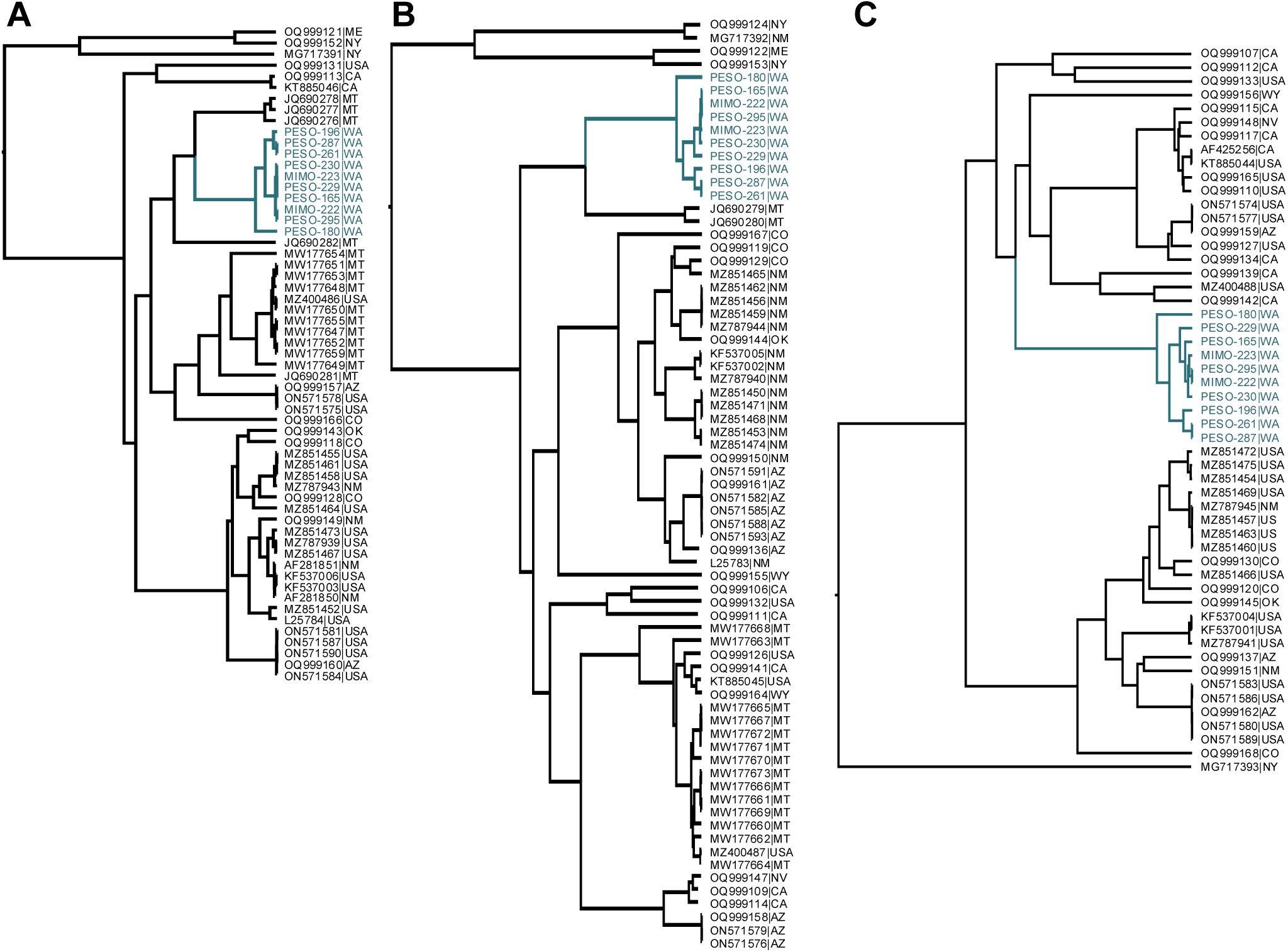
Phylogenetic reconstruction of the Washington samples of *Orthohantavirus sinnombreense* (SNV) in the context of viral diversity in the United States for the (A) S-segment, (B) M-segment, and (C) L-segment. The clade grouping the Washington samples generated in this study is highlighted in blue; MIMO indicates sequences from *Microtus montanus* and PESO indicates sequences from *Peromyscus sonoriensis*.

Discrete phylogeography based on the S-segment tree with the same MCC parameters(15) inferred introduction into Washington from Montana circa 1915 (HPD95%: 1873 – 1982), followed by local diversification (Figure 3A). Bayesian Stochastic Search Variable Selection analysis(15) shows low Bayes Factor support for Montana-Washington movement, suggesting unsampled intermediates and illustrating the need for continued genomic surveillance in the Northwestern United States.

**Figure 3.**
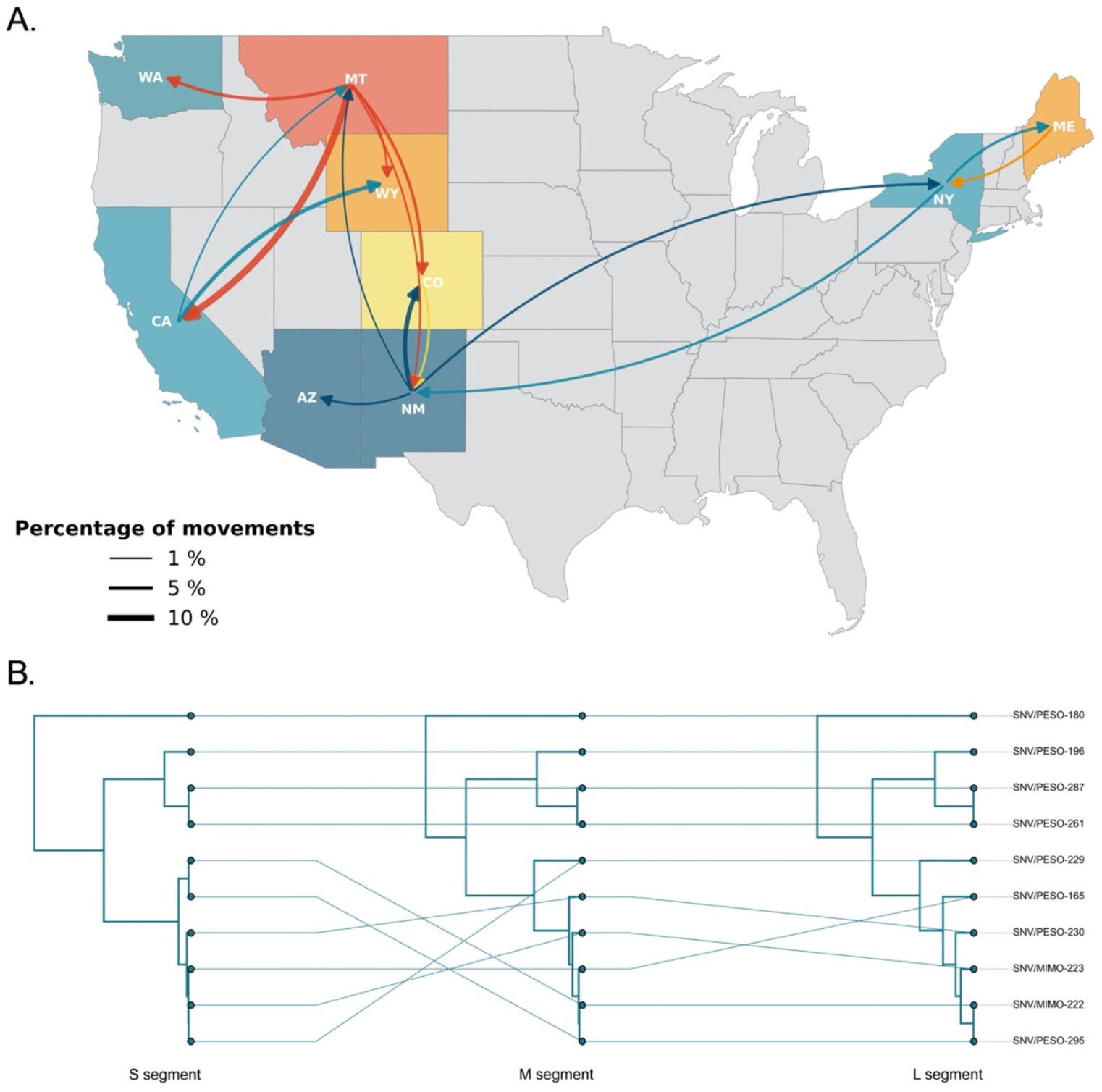
Evolutionary dynamics of SNV showing (A) phylogeographic reconstruction of movements between discrete states using Markov Jumps, with curve weight reflecting Bayes Factor support and (B) tanglegram showing topology changes across segment trees, where connecting lines link variants and major shifts indicate reassortment.

Topological discordance across viral segments supports viral reassortment in the local rodent population (Figure 3B). Within the Palouse clade, the montane vole (*Microtus montanus*, MIMO) derived variants shifted position across segment trees, alternately clustering with or diverging from the local western deer mouse-derived variants, further supporting reassortment and suggesting cross-species transmission between montane voles and western deer mice in the Northwestern United States (Figure 3B).

## Conclusions

We report on the first SNV genome sequences recovered from the Northwestern United States. Our findings highlight viral reassortment among sympatric rodent hosts of SNV, underscoring the complexity of SNV evolution and maintenance in rodents.

Understanding how multiple hosts contribute to these reassortment events will be crucial for clarifying transmission dynamics in wild rodents and anticipating risks of zoonotic emergence.

## Acknowledgements

This publication was made possible by cooperative agreement CDC-RFA-FT-23-0069 from the CDC’s Center for Forecasting and Outbreak Analytics. SNS, RR, and JH received support from US National Science Foundation (NSF DBI 2515340). The contents are solely the responsibility of the authors and do not necessarily represent the official views of the National Science Foundation or the Centers for Disease Control and Prevention. We thank Dr. Steven Bradfute for helpful discussions and guidance on recovering full viral genome sequences and the many field technicians who supported collections in the summer of 2023.

